# Mutational Analysis of Human Norovirus VP2 Elucidates Critical Molecular Interactions for Virus Assembly

**DOI:** 10.1101/2025.08.23.671901

**Authors:** Janam J. Dave, Sue E. Crawford, Robert L. Atmar, Khalil Ettayebi, B.V. Venkataram Prasad, Mary K. Estes

## Abstract

Human noroviruses (**HuNoV**s) are the leading cause of viral gastroenteritis with ≥80% of infections caused by the GII genogroup. HuNoVs are non-enveloped, with an icosahedral capsid composed of 90 dimers of the major capsid protein VP1, which encloses a minor structural protein, VP2, and a VPg-linked positive sense ssRNA genome. Although the atomic structure of the icosahedral capsid formed by VP1 is well characterized using crystallography and cryo-electron microscopy analyses of HuNoV virus-like particles (VLPs), the structures and the localization of VP2 and VPg inside the capsid, how they are incorporated into the capsid, and whether this process requires interactions between them remain unresolved. Herein, we show VP2 is the molecular bridge for assembly of particles containing VP1, VP2 and VPg. We used deletion constructs and mutational analyses, guided by bioinformatic analyses, to determine the interaction site on VP2 for VP1 of the pandemic-causing GII.4 Sydney HuNoV. GII.4 HuNoV VP2 contains a unique insertion site at amino acids (AAs) 43-53, relative to VP2s of other GII HuNoV genotypes. We identified AA residues 40-43 on VP2 are required for interaction with VP1; mutation of VP2 AA 40-43 abrogates VP2 encapsidation. Computational analyses predicted VP2 has a highly conserved N-terminal α-helical domain and an intrinsically disordered C-terminal domain that exhibits significant sequence diversity. We identified VP2, not VP1, uniquely binds VPg; the VP2 C-terminal domain is sufficient to interact with VPg. These findings reveal domain-specific functions of VP2 that are essential for coordinating capsid protein interactions for HuNoV assembly.

**Importance:** Human noroviruses (HuNoVs) are the leading cause of epidemic and sporadic gastroenteritis in all age groups worldwide. Yet, we currently lack vaccines or therapeutics for these pathogens. Knowledge about HuNoV biology is limited, including the fundamental mechanisms governing particle assembly. Modern structural techniques have not resolved the complete structure of pandemic GII.4 norovirus that includes the localization of the interior capsid proteins VP2 and VPg. Furthermore, VP2’s functional role(s) during infection remains obscure. Studies of feline and murine caliciviruses show VP2 may be involved in delivering the viral genome into cells, suggesting it synergizes with VP1 and VPg. We identify a motif on the N-terminal α-helical domain of VP2, adjacent to a unique insertion site, that is essential for interaction with the major capsid protein VP1. We show VP2 uniquely binds the translation initiation protein, VPg, via its disordered C-terminus. These findings reveal principles of HuNoV capsid protein interactions and highlight VP2 as a bridge facilitating capsid assembly.

## Introduction

Caliciviruses comprise a family of 11 genera and are non-enveloped, positive-sense RNA viruses that cause highly infectious diseases ranging from pandemic gastroenteritis in humans to hemorrhagic death in rabbits (1). Human noroviruses (HuNoVs) cause up to 680 million cases annually worldwide, resulting in 200,000 or more deaths globally and an estimated $60 billion economic burden each year (2–4). Although there was a recent surge in cases reported to be caused by GII.17 strains (5), most HuNoV cases result from infections with the globally-dominant GII.4 HuNoV (5–7). Despite their clinical importance, we lack antiviral therapies and vaccines for HuNoV (8). One barrier to the development of HuNoV antivirals is the lack of a fundamental mechanistic understanding of virus biology, including particle assembly, underscoring the importance of the continued study of this pathogen.

The molecular cloning of the GI.1 Norwalk virus (NV), the prototype HuNoV, revealed the HuNoV genomic RNA is organized into three open-reading frames (ORF1 through ORF3 (1, 9, 10). ORF1 encodes a polyprotein, which is autocatalytically processed by the virus-encoded cysteine 3C-like protease into six nonstructural proteins (11, 12). For HuNoVs, separate ORFs expressed from a subgenomic RNA encode the major and minor structural proteins that form the capsid. ORF2 encodes the major capsid protein, VP1, and a translational frameshift leads to expression of ORF3, which encodes the minor capsid protein, VP2 (10, 13). VP1 with or without VP2 can self-assemble to form VLPs (14, 15). High resolution structural characterization of these particles using X-ray crystallography or electron cryo-microscopy (cryo-EM) has revealed that these VLPs can adopt T=1, T=3, or T=4 icosahedral symmetry (16–19). However, the structures of native virions including murine norovirus (MNV), feline calicivirus (FCV), and San Miguel Sea lion virus determined using X-ray crystallography or cryo-electron microscopy show that their capsids exhibit T=3 icosahedral symmetry formed by 90 dimers of VP1 with a diameter of ∼40 nm, which is consistent with the diameter observed in the EM images for HuNoV virions (20, 21). Each VP1 subunit has a modular domain organization consisting of an internal N-terminal arm (NTA) and two distinct domains termed shell (S) and protruding (P) domains, separated by a flexible hinge (16, 17). Early studies showed that HuNoV VP1 and VP2 associate to form the virion, and VP2 is positioned inside the capsid during assembly by interacting with the VP1 S domain (13, 22). To date, structural techniques have failed to resolve HuNoV VP2, leaving the region of VP2 responsible for interaction with VP1 undefined.

There is little known about the functional role of VP2. Studies of FCV VP2 show it is required for production of infectious particles, underscoring its importance (23). Cryo-EM analyses of FCV in complex with its receptor, fJAM-A, show the capsid undergoes a conformational change resulting in the extrusion of a portal-like oligomeric structure, formed by the α-helical N-terminal region of VP2, which may be critical for delivering the viral genome into cells (24). Recent studies suggest FCV VP2 punctures a hole in the endosomal membrane to release the viral genome into the cytoplasm (25). In addition to VP2, all members of the *Caliciviridae* family contain VPg that is covalently attached to the ssRNA genome (1). In HuNoVs, VPg is required for RNA infectivity and genome replication (26). Whether VPg interacts with VP1 or VP2, or both proteins, during encapsidation is unknown. To address these knowledge gaps, we used truncations and mutational profiling with co-immunoprecipitation western blot analyses to identify the VP2 interaction motif for VP1 and to evaluate whether VP1 or VP2 recruits VPg during particle formation.

## Results

### VP2 consists of a conserved alpha helical domain and an intrinsically disordered domain

There are currently ten recognized norovirus genogroups (GI–GX) and two tentative groups that are further subdivided into 48 genotypes (6). HuNoVs in GI, GII, GIV, GVIII, and GIX genogroups, are genetically diverse, and sequence divergence is variable for each protein (6, 27). In GII viruses, VP2 and nonstructural protein 4 (NS4 or P22) exhibit the highest sequence diversity of any HuNoV proteins (28). Indeed, sequence alignment of representative strains from each HuNoV genogroup highlights evolutionary trends in VP2 (**Fig. 1**). HuNoV VP2s vary significantly in length. For example, GI.1 HuNoV VP2 is 212 AA, whereas those for GII.3 and GII.4 HuNoV are 254 and 268 AA, respectively (**Fig. 1**). The increased length in GII.4 VP2 is partly due to unique amino acid insertions at positions 43 and several smaller insertions proximal to the C-terminus, making GII.4 virus VP2 the longest HuNoV VP2 protein.

**Fig 1.**
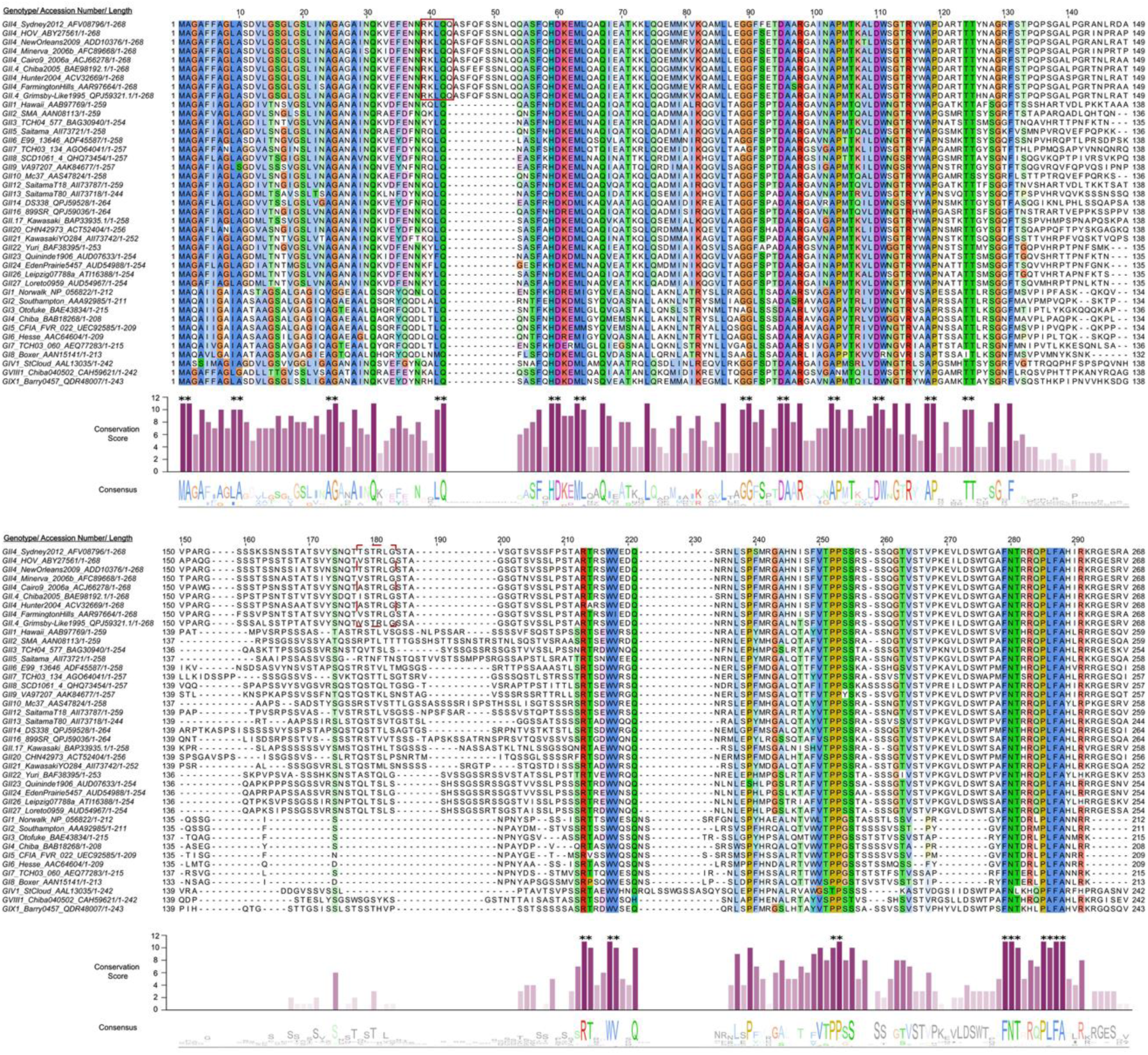
Amino acid sequence alignment of HuNoV VP2. Multiple sequence alignment of VP2 protein from representative HuNoV genogroups as designated by Chhabra *et al.* (6) was carried out in Jalview (29) using the ClustalO alignment algorithm (30). When a complete VP2 sequence was not found for a genogroup designated by Chhabra *et al*. (6), the closest corresponding VP2 sequence was substituted using a BLASTp search matching the respective VP1. **Top**: Multiple sequence alignment with residues colored based on ClustalX color scheme for biochemical properties and color saturation increased for ease of viewing. Blue = A, C, I, L, M, F, W, V, C; Red = K, R; Magenta = E, D; Polar = N, Q, S, T; Pink = Cysteine; Orange = Glycine; Yellow = Proline; Cyan = Histidine, Tyrosine. Representative members from each genotype are listed with NCBI accession number and AA length. HuNoVs are listed with genogroup and genotype, NCBI accession number and amino acid length of full-length protein. Numbers above residues indicate AA position based on sequence alignment. Red box indicates a region of high interest which was unique to GII.4 viruses and the focus of mutational analyses. Dashed red box at position 174-179 indicates a previously reported candidate region for a VP1 interaction site identified by yeast two hybrid studies (31, 32) that was tested by mutational analysis. **Middle**: Bar plot of VP2 AA conservation score (0–11) for each residue using JABAWS 2.2 (33); a score of 10 indicates a biochemically similar substitution in place of a consensus AA and a score of 11 indicates AAs with 100% conservation. Bar plot color intensity is a gradient scaled to conservation score, e.g. darker = more conserved. Asterisks are above consecutive AAs that scored 10 or 11 to highlight multiple consensus residue groups. The average consensus score in increments of 42 AA was used to compare conservation by region and the average scores used to designate regions were: < 3.5 = divergent, 5-7.5 unassigned, > 7.5 convergent. **Bottom**: Jalview-generated sequence logo depicting residue conservation across aligned VP2 sequences with residue conservation threshold set to ≥ 30%. Letter height reflects the degree of conservation at each position, with taller letters indicating higher conservation. Numbers at the end of each sequence alignment, bottom right, indicate actual length of protein, e.g. GII.4 Sydney 2012 is 268 AA.

In a multiple sequence alignment (MSA) of representative HuNoV VP2 sequences, we found the N-terminal half (AA 1-134) contains a higher density of conserved residues compared to the C-terminal half of the protein (AA 135-268) (**Fig. 1**). Using Jalview and the JABAWS consensus scoring system, multiple consecutive residues within the N-terminal region scored 10 or 11 (**Fig. 1, Middle,** indicated with asterisks above bar plot); a score of 10 is assigned for biochemically-like AA substitutions and a score of 11 represents complete conservation with no substitutions in the aligned sequences (33). Conversely, consensus scores in the C-terminal region dropped significantly, highlighting greater sequence variability (**Fig. 1**). There are fewer consensus residues in a majority of the C-terminal region except at aligned positions 213-214, 217-218, 252-253, and 279-290. The overall region spanning aligned positions 130-210 had an average conservation score below 3.5. Aligned positions 210-298 also had an average conservation score below 5.5, although within this region, positions 250-294 had an average score of 5.6. To further examine the structure-function relationship within HuNoV VP2, we used AlphaFold3 (34) to predict the VP2 structure. Alphafold3 predicts the VP2 N-terminal region is an α-helix (approximately residues 1–106), followed by a disordered C-terminal region (residues ∼123–268, the exact positions vary by genogroup) (**Fig. 2A, Fig. S.1**). For GII.4 Sydney VP2, the AlphaFold3 prediction has high confidence in the N-terminal α-helical region (predicted local distance difference test, plDDT >90, dark blue), transitioning to moderate confidence in the central region (plDDT 70– 90, light blue-yellow), and to low confidence (plDDT <50, orange) in the C-terminal domain (**Fig. 2B**). This sharp decline in plDDT score aligns with disorder predictions via IUPRED3, supporting a majority of the C-terminus being intrinsically disordered (35) (**Fig. 2C**). Conservation patterns mirror AlphaFold3 and IUPRED3 structural predictions, suggesting evolutionary constraints imposed on the α-helical domain. Despite high sequence diversity between genogroups, AlphaFold3 predicts HuNoV VP2 secondary structure is largely conserved, all representative genogroups encode an N-terminal α-helix and a variable C-terminal disordered region (**Fig. S1**). Overall, bioinformatic analyses and structural predictions indicate duality in VP2 architecture and sequence, where the N-terminal α-helical region is composed of highly conserved residues and the C-terminal region is mostly disordered (**Fig. 2D**).

**Fig. 2.**
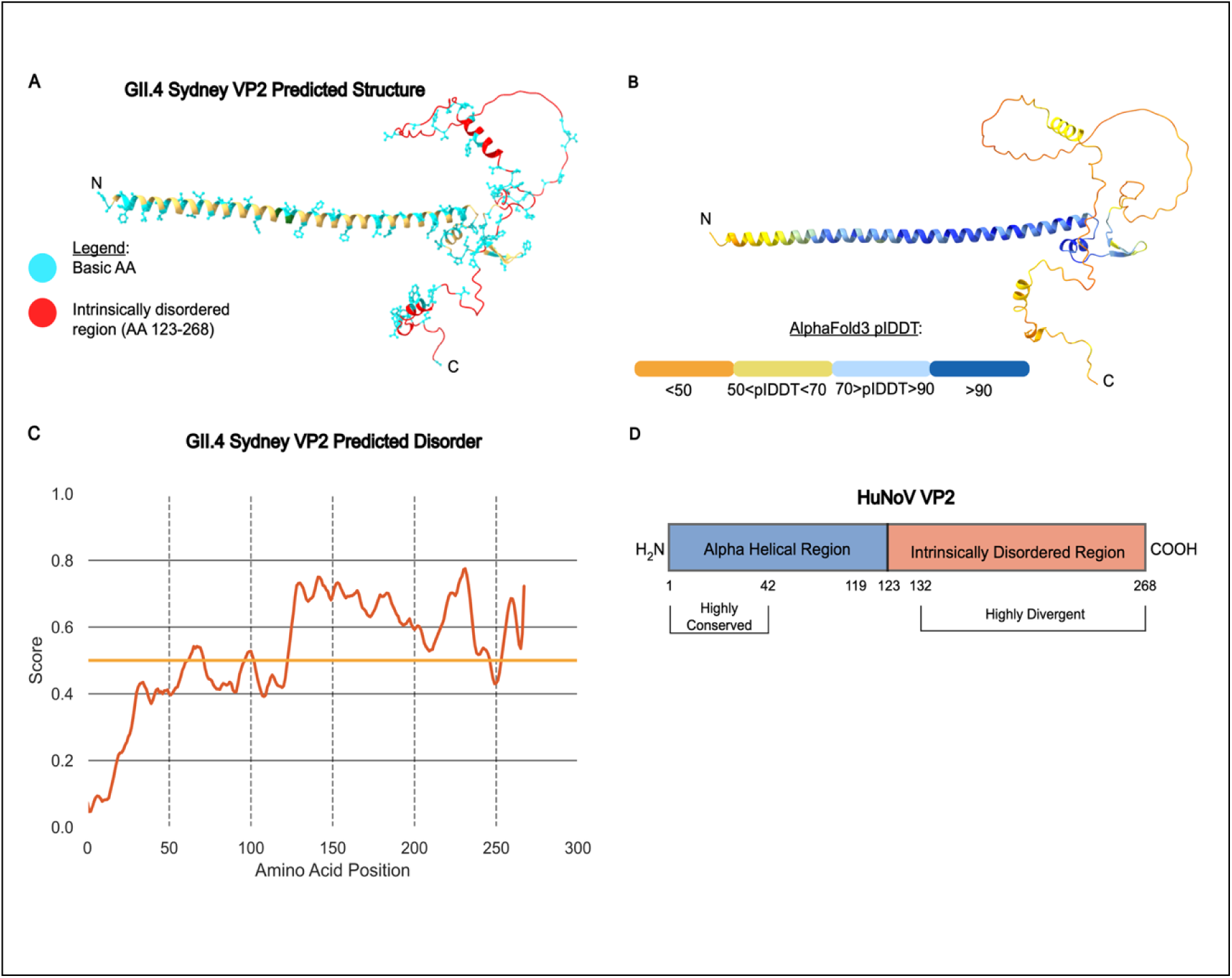
Multi-parameter analysis of HuNoV VP2 predicted structure, biochemistry and conservation. (**A** and **B**) AlphaFold3 predicted structures of GII.4 Sydney VP2. (**A**) Positions of basic residues and side chain (cyan) and its intrinsically disordered region (red) are highlighted, remaining regions of protein are colored gold and depict secondary structure in a ribbon model. (**B**) Predicted structure scored by plDDT scaled 0-100. (**C**) IUPred3-predicted intrinsic disorder score of HuNoV VP2 (disorder score cut-off of 0.5, long disorder, ≥ 30 consecutive residues). (**D**) Schematic summary and model of HuNoV VP2 structure and conservation. AA positions specific to GII.4 Sydney VP2 pandemic HuNoV.

We observed several residues in VP2 between AA 37-43 that are distinct in the globally circulating and pandemic-causing GII.4 viruses compared to other HuNoVs (**Fig. 1**, red box at AA 37-43). Further sequence alignment analysis using the NCBI non-redundant BLASTp database homology search for pandemic-causing GII.4 Sydney VP2 yielded a top cluster containing 1373 GII virus protein sequences. Analysis of this cluster reinforced our observations from the representative genogroup VP2 multiple sequence alignment (**Fig. 1**), which highlighted more focused conservation of N-terminal residues (**Table S1**). In this cluster, residues 37-43 contained the highest density of multiple consecutive consensus residues, including all known pandemic GII.4 strains (**Table S1**). We also found AA 41-43 and 109-111 were unique as they contained three consecutive AA with 100% consensus residues, with upstream AA 39-40 nearly reaching 100% consensus except for 2 out of the 1374 sequences (**Table S1**). Therefore, we probed this N-terminal region, up to AA 50, for possible VP1 interaction sites using co-immunoprecipitation (co-IP) and western blot analysis of 3xFLAG-tagged wild-type or mutated VP2 and wild-type VP1 co-expressed in HEK293FT cells (**Fig. 3A**).

**Fig. 3.**
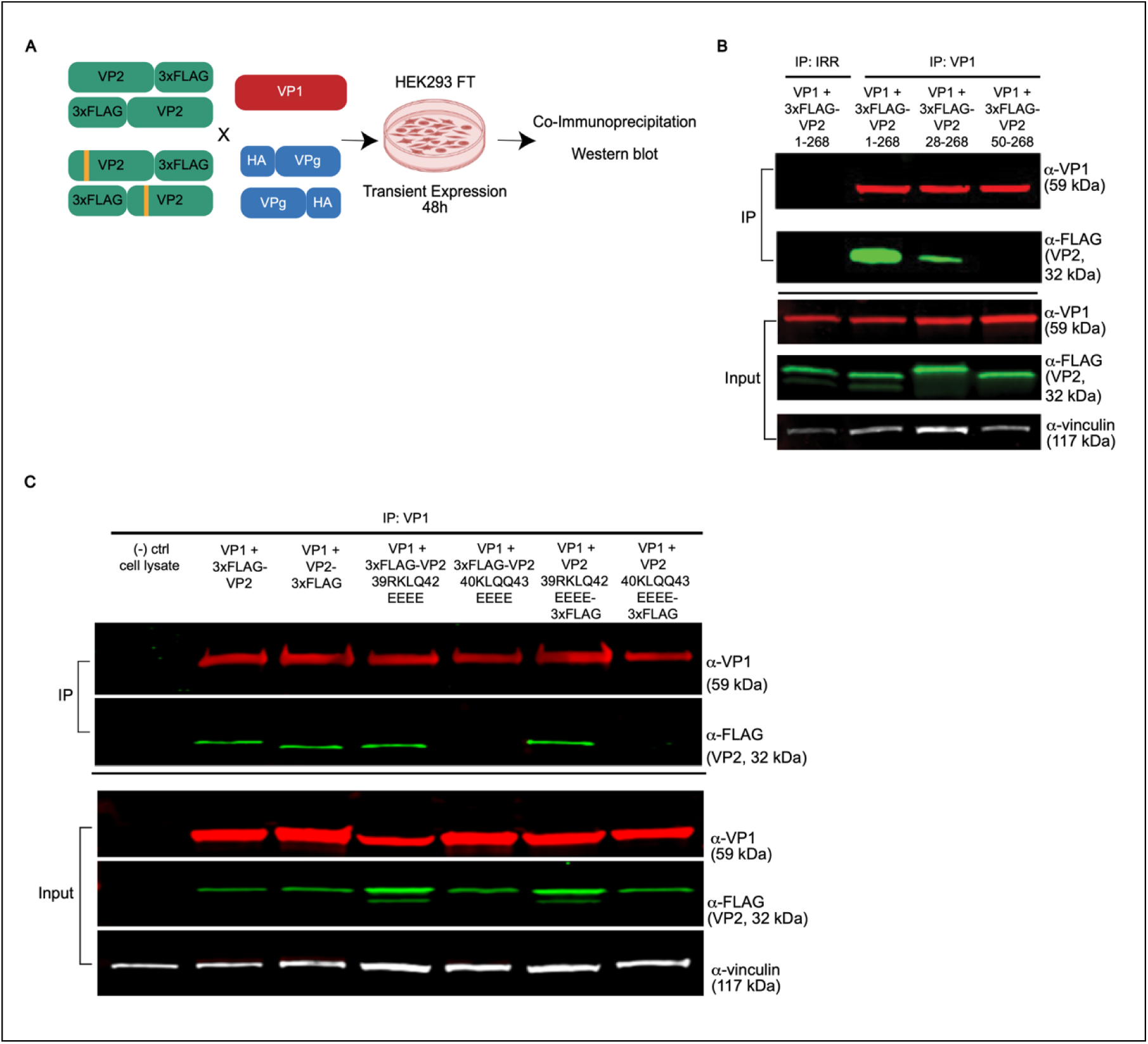
The highly conserved N terminal arm of VP2 encodes a putative VP1 interaction motif. (**A**) Experimental design for mutational analyses and recombinant protein co-expression in HEK 293FT cells to determine HuNoV capsid protein-protein interactions via co-IP western blot. (**B** and **C**) Co-IP western blot analysis of 3xFLAG VP2 interaction with WT VP1. The proteins indicated above each lane were co-expressed and immunoprecipitated. VP1 was pulled down using α-VP1 NV 23 mAb. Each protein was detected using the antibody indicated to the right of each immunoblot: VP1 was detected with α-VP1 rabbit pAb, VP2 detected via α-FLAG mouse M2 mAb, vinculin detected via rabbit α-vinculin rabbit mAb. Nitrocellulose membrane for western blot cut and only relevant data are shown. (**B**) VP1 interaction with 3xFLAG (N-terminus) full length VP2 (1–268), or VP2 truncated to AA 28-268 or 50-268. (**C**) VP1 expressed with either full length VP2 or VP2 mutants AA 39-42 (RKLQ) or 40-43 (KLQQ) substituted to 4x glutamate (EEEE) fused to a 3xFLAG tag at the N or C terminus were tested for interaction with VP1.

First, to test whether the VP1 interacting site is within the first 50 amino acids of VP2 or is further downstream, immunoprecipitations from cell lysates in which VP1 was co-expressed with full-length N-terminal 3xFLAG-VP2 (VP2 1-268), or truncated 3xFLAG-VP2 (VP2 28–268 or VP2 50–268) were performed. A VP1 monoclonal antibody (IP: VP1) was used to capture VP1 and co-IP interacting VP2. To control for nonspecific binding and validate the specificity of antibody capture in the co-IP, an irrelevant antibody control (IRR) matching the species IgG isotype was used (**Fig. 3B**, IP: IRR). Western blot analysis showed that VP1 interacted with full-length VP2 and VP2 28–268, whereas no interaction of VP2 50–268 with VP1 was detectable, indicating AA 28-50 are critical for VP1 binding (**Fig. 3B)**. Western blot analysis of co-IP inputs for VP1 and VP2 relative to the vinculin loading control confirmed successful expression of all constructs (**Fig. 3B**, input panel), indicating there are varying expression efficiencies for co-expression of VP1 and VP2 variants but a lack of VP2 interaction is not an artifact of a deficit in VP1 or VP2 expression levels.

To further dissect the conserved residues within the VP2 N-terminal region that interact with VP1, specific glutamate substitutions were introduced at the highly conserved residues 39–42 (RKLQ) and 40–43 (KLQQ). Separate 3xFLAG fusion tags at either the N- or C-termini (indicated as 3xFLAG-VP2 or VP2-3xFLAG, respectively) of recombinant full-length or mutated VP2 were tested to ensure results were not an artifact of fusion tag position since tag position can affect the protein folding and biochemistry (**Fig. 3C**). These mutants were co-expressed with VP1, and co-IP assays revealed mutation of VP2 AA 40KLQQ43 to glutamate (EEEE), but not 39RKLQ42 to glutamate (EEEE), abrogated VP1 interaction with VP2 with 3xFLAG tag at either the N- or C-termini (**Fig. 3C**). These findings indicate GII.4 HuNoV VP2 AA 40KLQQ43 as critical for interacting with VP1 and that the placement of the 3xFLAG tag at either terminus of VP2 did not influence this interaction.

### Complete mutation of AA 40-43 on VP2 is required to abrogate interaction with VP1

We next analyzed whether the entire KLQQ motif (AA 40–43) within the VP2 N-terminal domain was required for VP1 interaction. We observed that the dual “QQ” residues on AA 42-43 are largely conserved on GII viruses and particularly GII.4 HuNoVs (**Fig. 1**). AlphaFold3 structural modeling suggested a high degree of surface accessibility and proximity of VP2 QQ 42-43 that could potentially participate in hydrogen bonding with VP1, to mediate direct contact (**Fig. 4A**). Moreover, Q42 is conserved on all representative HuNoV VP2s but Q43 is specific to GII.4 viruses as the first AA in the specific inserted sequence and largely dispensable for non-GII viruses based on the HuNoV VP2 sequence alignment (**Fig. 1**). We initially substituted 40KLQQ43 to 40EEEE43 (**Fig. 3C**), which introduces a net negative charge at the 40K (positive) and 41LQQ43 (neutral) charged sequences. We also considered whether the loss of VP1 interaction was due to the introduction of a net negative charge on VP2’s interaction motif, so we mutated AA 40-43 to a set of uncharged residues, with a 4x alanine substitution. Overall, these observations led us to probe whether a dual 42QQ43 mutation to 2x glutamate or 2x alanine would achieve the same loss of interaction phenotype as the mutation of AA 40-43 to 4x glutamate or 4x alanine. Co-immunoprecipitation experiments with VP2 mutations at residues 42–43 (QQ) to alanine or glutamate showed VP2 still maintained its ability to bind VP1 (**Fig. 4B**), indicating that mutation of these two residues alone is insufficient to disrupt the interaction. Co-expression of VP1 with VP2 40AAAA43 containing a 3xFLAG tag at either the N- or C-terminus and co-IP assays revealed mutation of VP2 AA 40KLQQ43 to alanine (AAAA) abrogated VP1 interaction with VP2 with 3xFLAG tag at either the N- or C-termini (**Fig. 4B**). These findings indicate that substitution of VP2 40KLQQ43 to 40EEEE43, with a net negative charge, or to uncharged residues 40AAAA43, both abrogate the VP1-VP2 interaction and are not an artifact of a charge substitution.

**Fig. 4.**
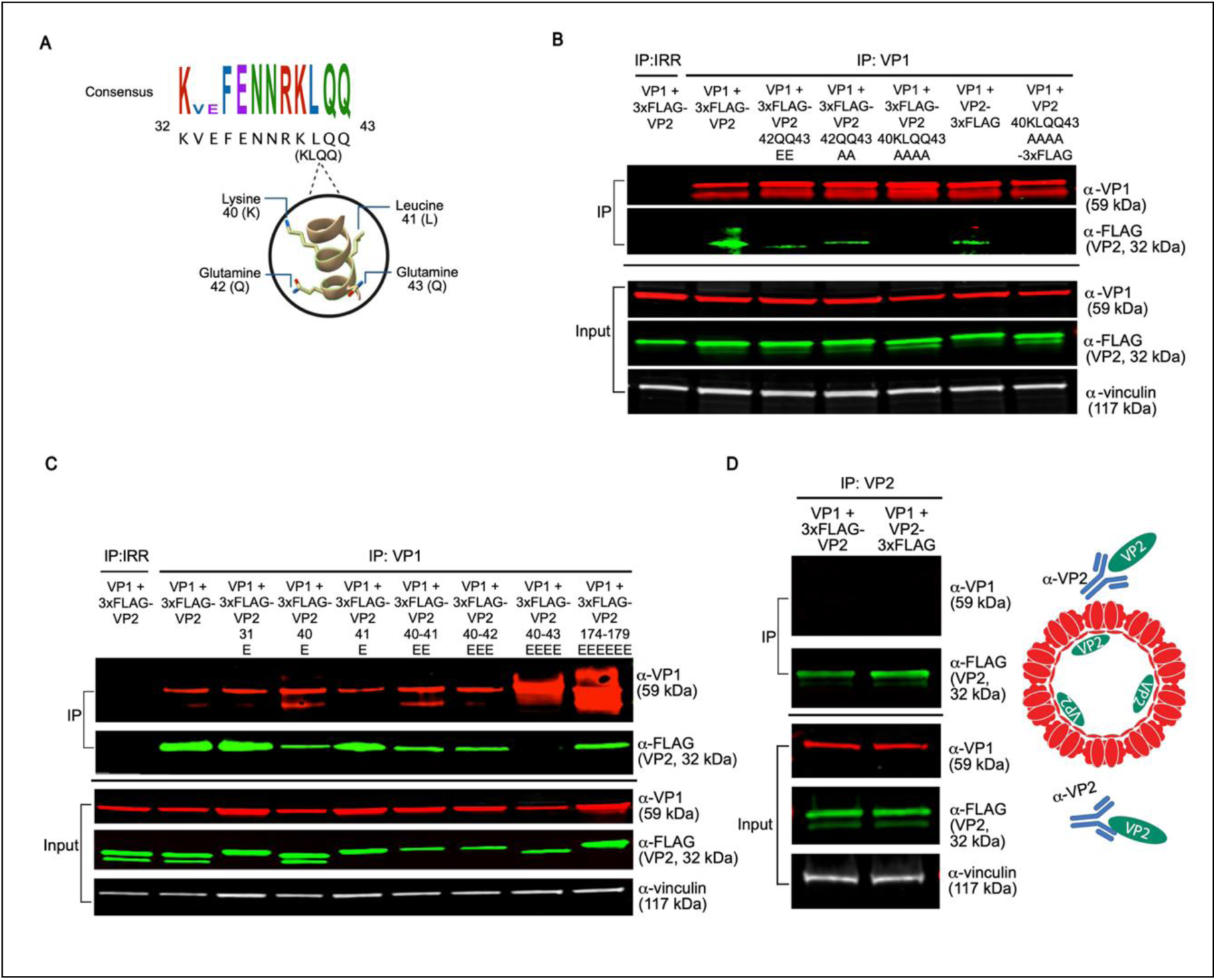
Mutation of AA 40-43 on VP2 is required to abrogate a VP1-VP2 interaction. (**A**) Sequence logo and AlphaFold 3 predicted structure of the amino acids of interest flanking the highly conserved LQ patch (AA 41-42) on HuNoV VP2’s VP1 interaction motif (AA 40-43) visualizing the AA position along its outer periodicity, 50<plDDT<70. (**B - D**) Western blot analysis of co-IP experiments with mutational scanning of putative VP1 interaction motif residues on VP2 (AA 40-43) and partial mutations on VP2 as described in Fig. 3B**-3C**. An N-terminal 3xFLAG fusion tag was used on VP2 unless otherwise indicated. (**B**) Western blot analysis of VP2-VP1 interaction with the indicated VP2 mutants. (**C**) Western blot analysis of VP2-VP1 interaction with single and consecutive mutations of AA 40-43 to glutamine. (**D**, **Left**) Western blot analysis of VP2-VP1 interaction with VP2 reciprocal immunoprecipitation in non-denaturing conditions (SDS free). (**D**, **Right**) Schematic of VP2 reciprocal IP results, indicating unbound cytosolic VP2 can be pulled down, however, the encapsulated VP2 is inaccessible to antibody targeting.

Extending our mutational analysis further, we generated additional single and consecutive glutamic acid mutants within the KLQQ motif to test whether a smaller set of residues within AA 40-43 was critical for interaction with VP1. We also tested VP2 mutant Q31E, another 100% conserved AA between AA 28-50 (**Fig. 1**). Unlike mutation of VP2 AA 40-43 to 4xE, mutation of a single residue, either Q31E, K40E or L41E, did not disrupt the VP2-VP1 interaction, nor did consecutive mutations (K40+L41 to EE) or triple mutations (K40+L41+Q42 to EEE) (**Fig. 4C**). We also examined a distal VP1 interaction motif candidate on VP2’s C-terminus, which was located at a region following another possible divergent insertion site relative to GII.4 VP2. We evaluated AA 174-179, as reports using yeast 2-hybrid screening proposed this C-terminal patch is critical for VP1 interaction GII VP2 (31, 32). This putative motif on GII.4 VP2 (residues 174TSTRLG179, refers to native AA positions not Fig. 1 sequence alignment AA positions) contains a highly conserved residue, serine 175, and is also between a patch of AA that GII.4 VP2 is missing but is present in multiple GII viruses including GII.1, GII.2, GII.12, GII.14, GII.16, GII.17 (**Fig. 1**). Co-IP of VP1 co-expressed with VP2 174EEEEEE179 resulted in VP1 interaction with the mutated VP2, indicating that this C-terminal patch is not critical for GII.4 VP1 interaction (**Fig. 4C**).

Recombinant expression of VP1 alone results in self-assembled homo-oligomers that form VLPs (15). To examine whether the VP2 in these assays is encapsidated into particles, we performed reciprocal immunoprecipitation targeting VP2, instead of VP1, under non-denaturing conditions (**Fig. 4D**, left). 3xFLAG-VP2 and VP2-3xFLAG were successfully co-immunoprecipitated when pulling down VP1, but a VP2-VP1 complex was not co-immunoprecipitated when pulling down VP2, indicating a fraction of VP2 is cytosolic and accessible for pull down (**Fig. 4D**). In addition, there is another VP2 fraction bound to VP1 and internalized inside a VLP that is only targeted with VP1 co-IP (**Fig. 4B-C**). Together, these findings reveal a composite of electrostatic and hydrophobic residues, spanning AA 40-43, on the highly conserved, α-helical domain of GII.4 Sydney VP2 critical for VP1 interaction.

### HuNoV VP2 is the molecular bridge facilitating particle assembly

Given that three proteins make up the HuNoV capsid, VP1, VP2 and VPg, we next investigated how this heteromeric complex forms. We speculated that the predicted intrinsically disordered region (IDR) on VP2 might interact with VPg, as this protein also exhibits IDRs that potentially could be critical to bind multiple partners (12, 36–38). In addition, we hypothesized that VP2 would interact with VPg given their internalized positions in the capsid and the limited surface area on VP1 for further internal capsid protein interactions (17). Co-IP western blot analysis of 3xFLAG VP2 or VP2 40EEEE43 co-expressed with HA-VPg or VPg-HA confirmed VP2 interacts with VPg irrespective of HA tag position (**Fig. 5A**, left) or VP2 mutation. Reciprocal IP targeting VPg also pulls down 3xFLAG-VP2 or VP2-3xFLAG, regardless of tag position or VP2 mutation, confirming that VP2 AA 40-43 are not essential for VPg binding and that VP2 interacts with VPg (**Fig. 5A**, right). We also co-expressed VP1 and HA-VPg or VPg-HA to test whether VP1 could bind VPg. Co-IP with either the VP1 antibody or HA antibody to pull down the HA-tagged VPg and western blot analysis showed VP1 and VPg do not interact (**Fig. 5B**). We next examined whether the putative α-helical domain (AA 1-106) or intrinsically disordered domain (AA 106-268) of VP2 is sufficient to pull down VPg. We found VP2 truncated to just its N-terminal domain (predicted by AlphaFold3, AA 1-106) did not pull down VPg but, like full length VP2 AA 1-268, the C-terminal IDR (predicted by AlphaFold3 and IUPRED3, AA 106-268) was sufficient to interact with VPg (**Fig. 5C**). These results show VP1 only interacts with VP2, but VP2 interacts with both VP1 and VPg (**Fig. 5D**).

**Fig. 5.**
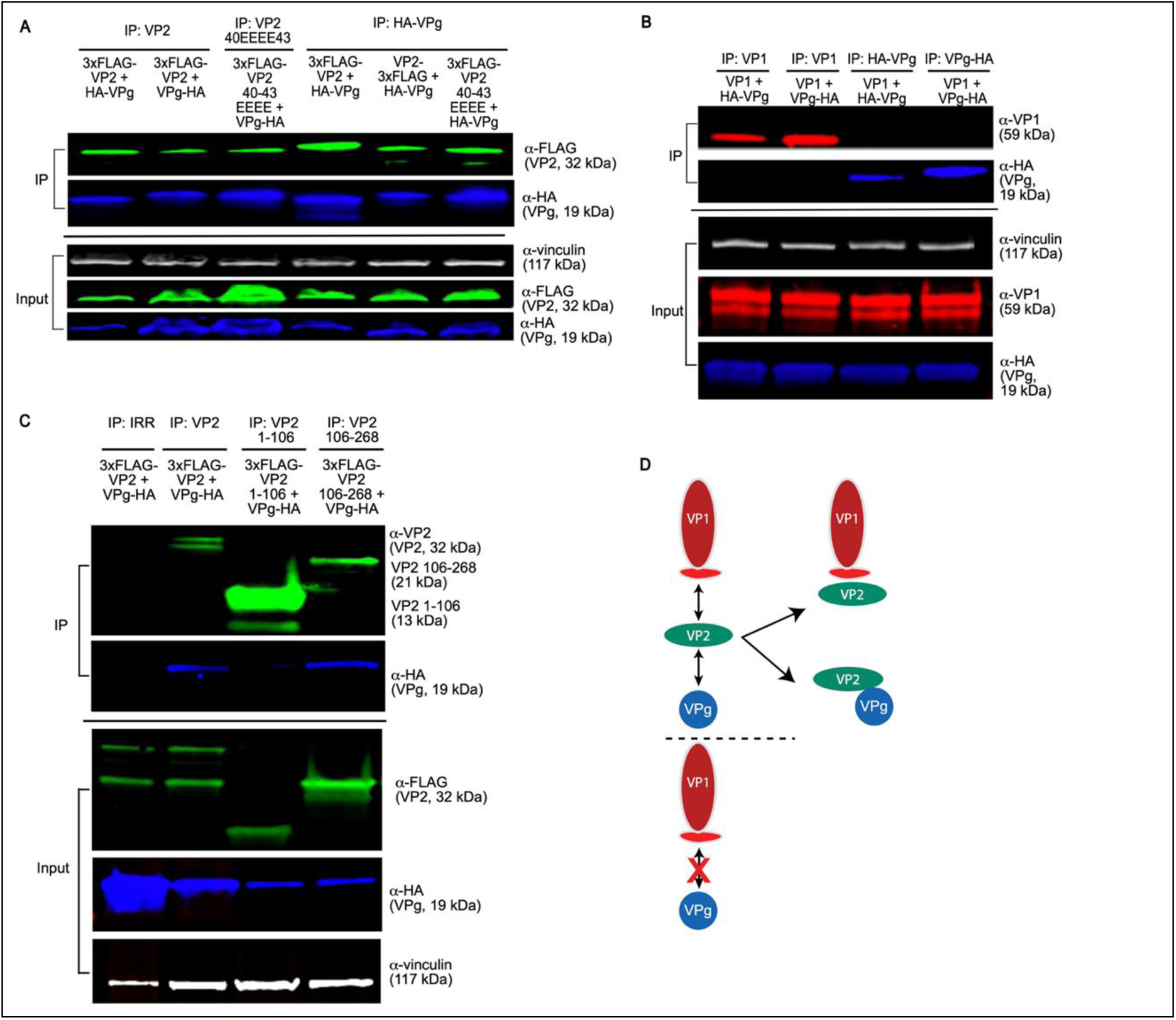
HuNoV VP2 is the only capsid protein that can interact with both VP1 and VPg capsid proteins. (**A**) Western blot analysis of 3x-FLAG VP2 co-expressed and co-immunoprecipitated with HA-VPg or VPg-HA (fusion tag placed at either N or C terminus, respectively). 3x-FLAG VP2 was pulled down using α-FLAG M2 mAb. VPg was detected via α-HA rabbit pAb and VP2 detected via α-FLAG M2 mAb, vinculin detected via rabbit α-vinculin mAb. Nitrocellulose membrane for western blot cut corresponding to virus protein size when probing IP samples and when probing for overlapping animal species, e.g. rabbit α-vinculin and rabbit α-HA. (**B**) Western blot analysis of VP1 co-expressed and co-immunoprecipitated with HA-VPg or VPg-HA. HuNoV VP1 pulled down using NV 23 α-VP1 mouse mAb. VP1 was detected via α-HoV rabbit pAb, and VPg detected as in **Fig. 5A**. (**C**) Western blot analysis of 3xFLAG VP2 full length or VP2AA 1-106 or VP2 AA 106-268) co-expressed and co-immunoprecipitated with VPg, proteins detected as in **Fig. 5A** above except VP2 IP fraction detected via VP2 rabbit pAb. (**D**) Schematic summary of capsid protein interactions assessed by IP and confirmed by reciprocal IP indicating VP2 can interact with VP1 and VPg but VP1 and VPg do not interact.

Since we observed VP2 could interact with either VP1 or VPg, we next considered whether a VP2 interaction with VP1 or VPg would preclude an interaction with its other counterpart. We tested this possibility for three reasons: i) all three proteins are known to make up the virus capsid, ii) given our results that identified a VP1 interaction motif on the N-terminus of VP2 while the C-terminal region on VP2 was sufficient to interact with VPg, we hypothesized VP2 molecules could interact with both VP1 and VPg concurrently, and iii) the VPg in these studies is not bound to the viral genomic RNA, which might be required for a VP1-VP2-VPg complex to co-IP. Immunoprecipitation and western blot analysis of VP1 and VPg co-expressed with or without VP2 confirmed VP1 cannot pull down VPg in the absence of VP2 (**Fig. 6A**, left). However, when AA 40-43 on the VP1 interaction motif (VIM) on VP2 are intact, VP1 pulls down VP2 and VPg interacting as a complex (**Fig. 6A**, center). Furthermore, expression of VP2 VIM mutants, AA 40KLQQ43 (to alanine 40AAAA43, VIM Δ_1_ or to glutamate 40EEEE43, VIM Δ_2_) resulted in particles composed only of VP1, indicating VP2 is required to recruit VPg inside the assembling complex (**Fig. 6A**, right). These results elucidate the interactions between capsid proteins for HuNoV particle assembly, which is dependent on VP2 acting as molecular bridge that uniquely interacts with both VP1 and VPg (**Fig. 6B**).

**Fig. 6.**
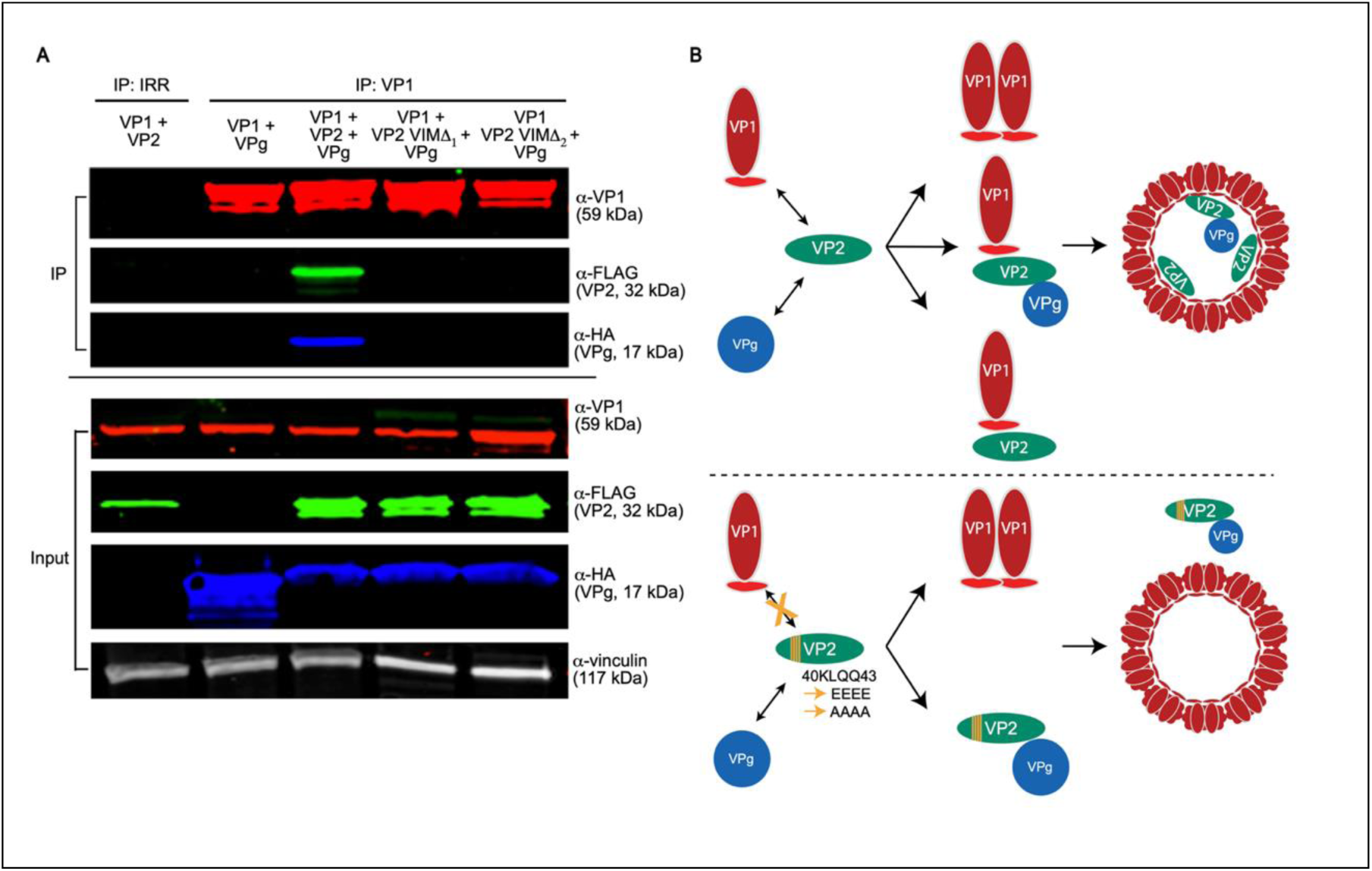
VP2 is required for formation of the VP1-VP2-VPg complex. (**A**) Western blot analysis of wild-type VP1, VP2-3xFLAG and HA-VPg interaction via co-expression and co-immunoprecipitation. The VIM on VP2 (AA 40-43 KLQQ) was mutated to 4x alanine (VIMΔ_1_ or 4x glutamate, VIMΔ_2_). The proteins indicated above each lane were co-expressed and co-immunoprecipitated using α-VP1 mAb. Each protein was detected using the antibody indicated to the right of each immunoblot as described in **Fig. 3B-3C** and **Fig. 5** above. (**B**) Model for HuNoV assembly indicating HuNoV VP2 interacts with VP1 and also recruits VPg inside the particle complex. A mutation on VP2 AA 40-43 disrupts particle assembly.

## Discussion

HuNoV capsid formation, in which the viral proteins self-assemble into an ordered architecture while overcoming host innate immunity, is a fascinating biological process that remains poorly understood. The mechanisms by which the internal capsid proteins, VP2 and VPg, interact with VP1 during encapsidation are lacking. In the present study, we addressed the fundamentals of internal capsid protein interactions and VP2’s role in assembly by using computational analyses to guide our biochemical investigations. We found HuNoV VP2 varies greatly in length and in sequence, due to a striking insertion at AA residues 43-53 unique to GII.4 VP2 and several C-terminal sites which contribute to GII.4 divergence (**Fig. 1**). It is also intriguing that GII.4 HuNoVs encode the longest VP2, given all six global HuNoV pandemics were caused by GII.4 HuNoVs.

To probe for a critical VP1 interaction motif or VIM on VP2, we focused on sequence conservation and looked closely at highly conserved AA motifs. We found a VIM on the highly conserved patch spanning AA 40-43 (**Fig. 4A-4C**) that is unique to GII.4 HuNoVs. These data suggest the GII.4 VP2 VIM we have identified could be genotype-specific. Previous studies have proposed VIMs on GI.1 VP2 (broadly on AA 108-152) expressed in Sf9 cells without any mutational analyses (13). In addition, putative VP1 interaction sites have also been reported in other GII viruses using yeast 2-hybrid screens; these studies included GII.2 (AA 167-178 and 184-186) (32) and GII.4 viruses (AA 131-160 and 171-180) (31, 39). When we mutated AA 174-179, to 4xE, which encodes the only highly conserved GII.4 residue, S175 in a “TSTRLG” motif (see red dotted box in Fig.1), VP2 interacted with VP1. It is difficult to draw parallels between these studies and our results since yeast 2-hybrid experiments involve fusing virus proteins with bulky host proteins (∼15-30 kDa) and forced nuclear localization, possibly limiting physiological relevance. We posit that our approach using mutational analyses and co-IP western blot studies with small fusion tags (1-3 kDa) in the HEK293FT human cell line is more representative in modeling VP1-VP2 interactions. Since the regions identified using yeast 2-hybrid screens are not conserved in the VP2s of a majority of HuNoVs from representative strains, and in some cases even GII.4 HuNoVs, it is also difficult to understand the structural and biochemical significance of a VP1-VP2 interaction at these putative sites. Yeast 2-hybrid screens may have identified ancillary interaction sites on VP2’s C-terminus, and our studies do not address this possibility.

Our initial studies identified that the internally located N-terminal region of VP1 in the prototype GI.1 and GII.4 HuNoV interacts with VP2, and mutation of a single isoleucine shell domain residue disrupts VP1’s ability to interact with VP2 (22). The discovery of a VP1 interaction motif, or VIM, on VP2 establishes a new understanding of the residues at the VP1-VP2 interface. This VIM (KLQQ AA 40-43) on VP2 lies within the N-terminal α-helical domain, predicted with high confidence by AlphaFold3, and its high conservation across many GII.4 HuNoVs suggest it is under functional constraint. More conclusively, mutagenesis experiments show that full disruption of all four VIM residues are required to abrogate VP1 binding, suggesting that the interaction is not mediated by a single residue but it requires a composite surface of complementary charge and hydrophobic interactions between VP1 and VP2. Our studies cannot determine whether the mutation of the VP2’s VIM that prevents VP1 interaction causes an indirect conformational change in another region of VP2. It is possible that our selected mutations obscure conformational epitopes or introduce conformational changes on VP2. However, structural predictions of VP2 with AA 40-43 mutations indicate the secondary structure of the alpha helical region is largely preserved, especially at AA 40-43. With mutation of AA 40-43 to 4x alanines or 4x glutamates, the predicted structure of the C-terminal IDR had some changes in orientation and folding but none seemed consequential (**Fig. S2**). However, we found chemically distinct substitutions on GII.4 VP2 (4x glutamate and 4x alanine) produced the same loss-of-function phenotype, suggesting these mutations directly disrupt a critical interacting surface. More likely, the N-terminal VIM is an essential and primary contact site for binding, and there may be multiple ancillary stabilizing sites elsewhere on VP2.

Our present study has limitations since it involves overexpression of virus proteins in HEK cells. Although co-IP studies are a widely accepted approach for dissecting protein-protein interactions, they do not directly provide structural confirmation or whether protein interactions are direct or indirect, where undefined bridging molecules might facilitate protein binding. Secondly, HEK cells do not encode the still unidentified HuNoV entry (26) coreceptor and so are not a fully permissive cell line for this virus. We have previously reported that transfection of the HuNoV genome into HEK cells leads to production of virus particles, showing that HEK cells have all required machinery to support particle assembly, but virus infection does not spread in HEK cells since they lack the entry receptor. Better understanding the nature of capsid protein interactions by obtaining an atomic structure of the VP1-VP2 complex using single particle reconstruction of purified VP1+VP2 VLPs may be helpful in further resolving these caveats. A putative VP1-VP2 stoichiometric ratio of 180 VP1 molecules to less than 10 VP2 molecules, was implied from the GII.4 HoV VLP cryo-EM and crystal structure (17). The structure of VP2 has not yet been resolved inside VLPs possibly because of the low copy numbers of VP2 per particle (17). Consistent with these predictions, our reciprocal IP studies indicate the ratio of VP1-VP2 interactions is such that VP1 clearly outnumbers VP2 in VP1-VP2 complexes, and an unbound pool of cytosolic VP2 is present in HEK cells overexpressing VP2 of which only a subset of VP2 binds VP1 (**Fig. 4D**).

Recent reports are advancing our knowledge of the potential roles of VP2 and its functional domains. VP2 is required to produce infectious murine norovirus and feline calicivirus (23, 40). FCV VP2 is involved in early events in the viral life cycle through interactions with the receptor fJAM-A and the endosomal membranes to facilitate delivery of viral RNA into the cell (24, 25). VP2’s RNA secondary structure along its N-terminus may contain regulatory elements that are required for productive infection, and its C-terminus is indispensable for animal virus infections (40). Although VP2 enhances particle stability and homogeneity (14), it has not been completely clear why VP2 is required for infectious particles. We found VP2, but not VP1, uniquely binds VPg and that its C-terminus is sufficient to do so (**Fig. 5A-5C**). These data also revealed that the C-terminal IDR of VP2 serves an essential function and highlight that VP2 is a mediator of morphogenesis because it links the outer VP1 shell to the internalized VPg (**Fig. 6A-6B**). Although there are major differences in FCV, MNV, and HuNoV VP2 sequences and length, the IDR region is a conserved feature of VP2, which we found was sufficient to pull down VPg in GII.4 HuNoV. We find the VP2-VPg interaction intriguing as it supports the involvement of VP2-VPg at both ends of the virus replication cycle, unpackaging the genome at the onset of infection and helping recruit VPg for particle assembly. The presence of extended IDRs in VP2 and VPg are hallmarks of viral proteins that undergo dynamic multivalent interactions and are used by viruses to induce liquid–liquid phase separation (LLPS) (41, 42), a mechanism HuNoV likely employs using multiple viral proteins to generate replication niches (43). Drawing from our results and recent reports, VP2 is possibly essential for producing infectious particles because it recruits VPg and mediates assembly, helps deliver the genome at the onset of infection, contains regulatory elements important during replication and packaging, and may serve as an RNA-binding protein. Whether virions lacking VP2 will be absent of VPg or have an intact viral genome should be explored in future studies. This question is more tractable in animal caliciviruses, which benefit from having a high-yield reverse genetics system that can be used to further dissect VP2-VPg interactions. Since VP2 is required for productive animal calicivirus infections, the conservation of VP2 secondary structure, particularly in the N-terminal region that contains the VP1-interacting motif, suggests that VP2 could mediate capsid assembly in many other caliciviruses. This is supported by the cryo-EM structure of FCV bound to its receptor, which shows an α-helical structure for the N-terminus of VP2 that is consistent with our AlphaFold3 predictions (24). In addition, the VP2 C-terminus may also recruit VPg in other caliciviruses; this idea is supported by recent studies showing the C-terminal region of VP2 is required for producing infectious murine norovirus particles (40). Our study reveals a central role for VP2 in HuNoV particle assembly. Together, our results enhance our understanding of VP2’s role in HuNoV capsid assembly and offer a new paradigm wherein VP2 potentially coordinates capsid morphogenesis by linking the outer VP1 shell and internal payload containing VPg.

## Materials and Methods

### Cloning of Recombinant Plasmids

HuNoV VP1, VP2 and VPg encoding mammalian plasmids were made using In-Fusion® snap assembly (Takara Bio, #638949). Individual cDNA synthesized fragments for VP1 and VPg were synthesized by Genscript, VP2 cDNA was synthesized by Genewiz. Fragments were cloned into the pCG backbone (Addgene, Plasmid#51476) using InFusion® assembly, fragment forward primer 5’-CTTAATTAAGCGACCATGGACTACAAGGATGACGATGACAAGGA-3’, reverse primer 5’-AAACAGTCGAGCTTATCCCCTCGCTTACGAATGTGAG-3’ and backbone forward primer 5’-TAAGCTCGACTGTTTAAACCTGCAG-3’ and reverse primer 5’-GGTCGCTTAATTAAGGATCCGAATTCG-3’. Truncated VP2 AA 28-268 or 50-268 were similarly cloned with InFusion® assembly using the 3xFLAG VP2 as a template. VP2 N- and C-terminal 3xFLAG-tagged protein glutamate and alanine substitution mutants were made using site-directed-mutagenesis by Epoch Life Science.

### Co-Expression

HEK293FT cells were transfected with recombinant plasmids using the TransIT-LT1 transfection reagent (Mirus Bio™□, #2300). Transfection efficiency and recombinant capsid protein expression were verified using the Zeiss Laser Scanning Microscope LSM 980 or Olympus epifluorescence IX73 (S016724) via immunofluorescence.

### Co-immunoprecipitation

Transfected cells were incubated for 48h post-transfection and lysed with IP Lysis buffer (Pierce, #87787, non-denaturing experiments) or RIPA buffer (Pierce, #89900) for 1h at 4°C with Benzonase (Millipore, #E1014) and protease inhibitor cocktail: cOmplete™□(Roche, #11873580001) with aprotinin (Sigma, #A1153), leupeptin (Sigma, #L2284), pepstatinA (Sigma, #P4265), PMSF, (Sigma #10837091001) and phosphatase inhibitor cocktail 2 and 3 (Sigma, 2 - #P5726, 3-#P0044). The clarified supernatants were incubated with the following antibodies for 2h at 4°C: VP1 pull downs used 40 – 80 µg of NV 23 mouse anti-VP1 monoclonal (44), VPg pull downs used 10 µg of rabbit anti-HA (CST, #A3724) and mouse anti-HA (Thermofisher #26183), 1h at 4°C with M2 mouse 36 µL of anti-FLAG affinity gel (Millipore, #A2220) and for irrelevant IP controls, the mouse anti-V5 IgG_1_ antibody was used (Thermofisher, #R960-25). For VP2-VPg and VP2-VP1 reciprocal pull downs, sodium dodecyl sulfate (SDS) was excluded from buffers, as it is strongly advised by the manufacturer to prevent M2 antibody denaturation.

For VP1 pull down with VP2 or VPg, the anti-VP1 antibody-lysate mixture was then incubated with protein A/G beads for 1h (Pierce, #88802) at 4°C. For VP1-VP2 IPs, the immunoprecipitate was then washed 2x with RIPA lysis buffer with SDS and NaCl (25 mM Tris-HCl pH 7.6, 500 mM NaCl, 1% NP-40, 1% sodium deoxycholate, 0.3% SDS) containing protease inhibitor cocktail and 1 mM MgCl_2_, 500 mM NaCl (wash 1) or 150 mM NaCl (wash 2), and benzonase to remove nucleic acids. For VP2-VPg or VP1-VPg pull downs, the immunoprecipitate was washed 2x with IP lysis buffer and 1x with PBS pH 7.4. For VP1-VP2-VPg IPs, the immunoprecipitate was washed 4x with: (wash 1) - 0.5 M NaCl, 25 mM Tris-HCl, 0.1% vol/vol NP40, (wash 2) – 0.15M NaCl, 1 mM EDTA, 0.01 M Tris-HCl, pH 7.4, 0.1 % vol/vol NP40, 0.3% wt/vol SDS, (wash 3) - 0.01 M Tris-HCl, 0.1% vol/vol NP40, (wash 4) PBS all at pH 7.4.

### Western blotting

Nitrocellulose membranes were cut at corresponding protein sizes then probed with the following primary antibodies: VP1 was detected via rabbit anti-HoV VP1 pAb (44), VP2 was detected via mouse anti-FLAG M2 mAb or VP2 anti-rabbit pAb, and VPg was detected via rabbit anti-HA mAb. Proteins were detected using the Li-Cor Odyssey Infrared imaging system and IRDye® 800CW and IRDye® 680RD infrared dyes conjugated donkey anti-rabbit, anti-mouse and secondary antibodies, purchased from Li-Cor Biosciences (Lincoln, NE). This method enables quantitative detection over a broad dynamic range (∼4–6 logs) with high sensitivity, detecting as little as 1–2 ng of target protein per band. Commercial antibodies were tested for the presence of pre-existing norovirus antibodies and specificity by western blot. Only antibodies found to be negative for HuNoV antibodies and high specificity were used in these studies. For detection of VP2 in experiments containing truncated VP2 pull down for VPg (**Fig. 5C**), VP2 was detected with an affinity purified rabbit anti-VP2 polyclonal antibody developed by inoculating rabbits with expressed and purified VP2 AA 49-231 (ABClonal). Animal sera were tested before inoculation by ELISA for VP2 and by western blot for GII.4 Sydney 2012 HuNoV VP1, VPg and 3xFLAG VP2 to verify the animals were not previously exposed to HuNoVs. Rabbit terminal bleed sera was affinity purified using a protein A/G capture column for anti-VP2 polyclonal antibodies and similarly verified for sensitivity and specificity and cross-reactivity to endogenous proteins as well as other HuNoV proteins. 3xFLAG-VP2 was used as a positive control to confirm the rabbit polyclonal detects the same VP2 band as the M2 anti-Flag mAb.

### Computational analysis

HuNoV sequences were retrieved from NIH NCBI and GenBank published sequences and visualized in Jalview. HuNoV genogroups designated as representative by Chhabra *et al.* (6), were used for multiple sequence alignment (**Fig. 1**). Eight GII.4 variants previously designated as globally circulating are included in this alignment, which also includes the 6 global pandemic causing GII.4 viruses. Finally, GII.4 HoV was included as we previously published the VP1 interaction site for VP2 for this variant and for the GI.1 Norwalk virus (22). Where a complete VP2 sequence could not be found for the representative genogroup, a substitute VP2 sequence was identified using a BLAST search for the closest matching NCBI entry based on the respective VP1 sequence (GII.4_Minerva, GII.4_Cairo, GII.4_Grimsby-Like, GII.17 Kawaski). An NCBI BLASTp non-redundant search was also performed to find the top cluster matching GII.4 Sydney VP2 sequence (NCBI AFV08796), which yielded 1373 sequences from multiple GII virus genotypes. Residue conservation cut off was set to 30%. IUPRED3 cutoff was set to 0.5 and disorder set to long disorder with no smoothing. Protein structure and molecular modeling or docking simulations were performed using AlphaFold3, HADDOCK, and ChimeraX to visualize, model and recolor molecules (45–47).

## Data Availability

Further supporting data can be found at https://github.com/JanamDave/source-code. The Github repository includes multiple sequence alignment (MSA) data for Fig. 1, MSA annotations and calculated data, 1374 sequences from the top GII.4 Sydney 2012 VP2 cluster from a blastP non-redundant database search and their respective annotated and calculated raw data summarized in Table S1. A list of updates to this repository will be available with contemporaneous updates and date of changes, if any, in its “Read Me” section.

## Acknowledgements

This research was supported by Public Health Service grants from the National Institutes of Health for grants P01 AI 057788 and U19 AI 116497 (to MKE), S10 OD028480 that supported purchasing the Zeiss Laser Scanning Microscope LSM 980 with Airyscan 2, and the Olympus epifluorescence IX73 (S016724). The authors would like to thank Dr. Sasirekha Ramani (BCM), Dr. Anish Ramakrishnan (BCM), Dr. Liya Hu (MD Anderson) and Dr. Wilhelm Salmen (University of Michigan) for insightful discussions, review, and aid in using structural programs.

## Author Contributions

J.J.D. contributed to conceptualization, methodology, investigation, formal analysis, writing – original draft, and writing – review and editing.

S.E.C. contributed to conceptualization, methodology, formal analysis, and writing – review and editing.

R.L.A. contributed to formal analysis, assisted with multiple sequence alignment development, and contributed to writing – review and editing.

K.E. contributed to methodology, investigation, and writing – review and editing.

B.V.V.P. contributed to conceptualization, methodology, and writing – review and editing.

M.K.E. served as supervisor and corresponding author, and contributed to conceptualization, formal analysis, writing – original draft, and writing – review and editing.

---

**Fig. S1.**
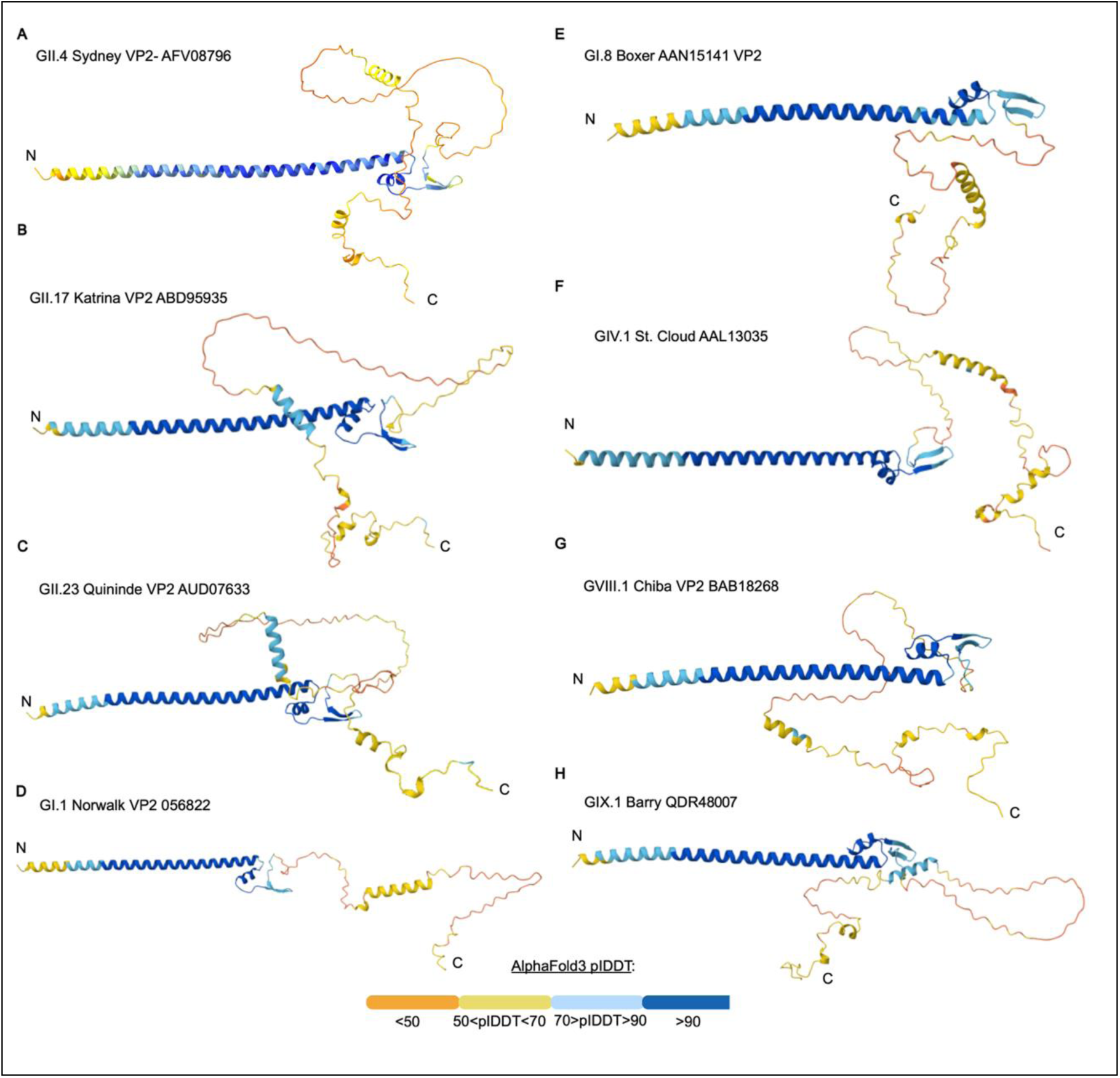
Alphafold3 predicted structures of HuNoV VP2 from representative genogroups. Sequences were selected from at least one representative genogroup, including (**A**) GII.4 Sydney (NCBI accession #AFV08756), (**B**) GII.17 Katrina (#ABD95935), (**C**) GII.23 Quininde (#AUD07633), (**D**) GI.1 Norwalk (#056822), (**E**) GI.8 Boxer (#AAN15141), (**F**) GIV.1 St. Cloud (#AAL13035), (**G**) GVIII.1 Chiba (#BAB18268), and (**H**) GIX.1 Barry (#QDR45007). For each VP2 model, the N- and C-termini are labeled. Structures are colored by plDDT confidence scores: blue (plDDT > 90, high confidence), cyan (70 < plDDT ≤ 90), yellow (50 < pLDDT ≤ 70), and orange (plDDT ≤50, low confidence). The predicted α-helical N-terminal domains are consistently modeled with high confidence, while the C-terminal regions are more variable in length and secondary structure and often predicted to be disordered. Representative genogroups based on designation by *Chhabra et al*. (6).

**Fig. S2.**
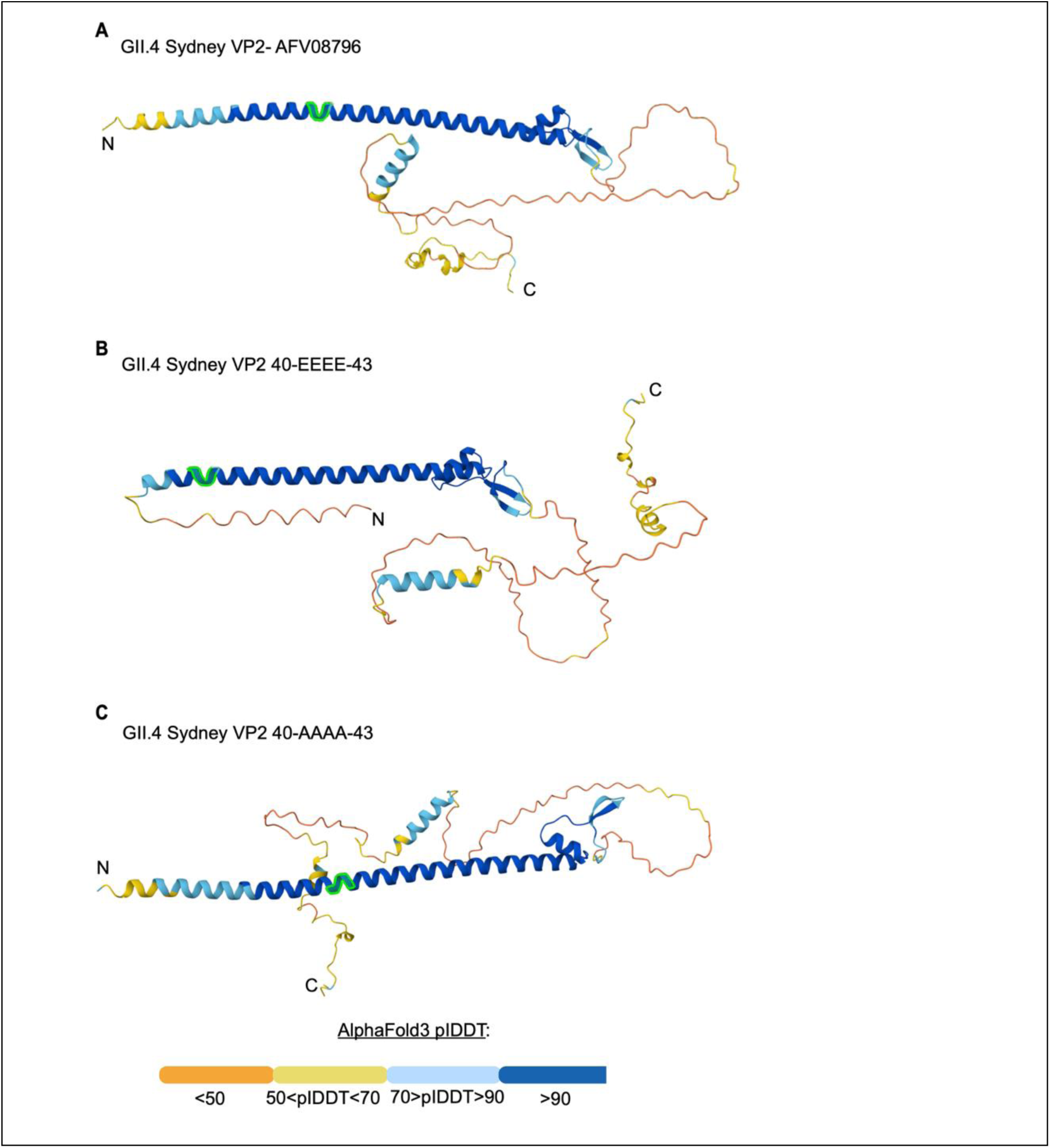
Alphafold3 predicted structures of wild-type HuNoV VP2 or recombinant VP2 mutated at AA 40-43. (**A-C**) AlphaFold3 predicted structure of VP2 with N- and C-termini labeled. AA 40-43 are highlighted (green) in each predicted structure. (A) Predicted structure of wild-type VP2. (**B**) Predicted VP2 structure with AA 40-43 mutated to 4x glutamate and (**C**) AA 40-43 mutated to 4x alanine. The α-helical N-terminal domains at mutation site are consistently modeled with high confidence, while the C-terminal regions are more variable in length and secondary structure.

